# Vaginal bacteria-derived extracellular vesicles diffuse through human cervicovaginal mucus to enable bacterial signaling to upper female reproductive tract tissues

**DOI:** 10.1101/2025.04.28.651063

**Authors:** Darby Steinman, Alyssa Petersen, Yasmi Chibber, Hannah C. Zierden

## Abstract

The composition of the vaginal microenvironment has significant implications for gynecologic and obstetric outcomes. Where a *Lactobacillus*-dominated microenvironment is typically considered optimal, a polymicrobial environment is associated with increased risk for female reproductive diseases. Recent work has examined bacteria-derived extracellular vesicles (bEVs) as an important mode of microbe-host communication in the female reproductive tract, with bEVs exhibiting unique species- and strain-level functions that may influence women’s health outcomes. However, in order to communicate with female reproductive tissues, bEVs must be able to penetrate the protective cervicovaginal mucus barrier. As the first line of defense against bacteria and pathogens, cervicovaginal mucus protects against infection in the female reproductive tract through steric, hydrophobic, and electrostatic interactions with foreign pathogens. Here, we hypothesize that the physical properties of bacteria-derived extracellular vesicles enable their mobility through cervicovaginal mucus and permit interactions with upper female reproductive tract tissues. We demonstrate that the barrier properties of mucus allow increased diffusion of bEVs, compared to whole bacteria. We evaluate the uptake of bEVs by, and the resulting effects on, human vaginal epithelial, endometrial, and placental cells, highlighting potential mechanisms of action by which vaginal dysbiosis contributes to gynecologic and obstetric diseases. Our work demonstrates the ability of bEVs to mediate female reproductive diseases and highlights their potential as therapeutic modalities for treating dysbiosis and dysbiosis-associated diseases in the female reproductive tract.

## Introduction

The vaginal microbiome significantly influences gynecologic and obstetric outcomes. Unlike most microbial communities in the human body, an optimal vaginal microbiome is one dominated by lactobacilli, which produce antimicrobial, antiviral, and antifungal agents that protect the local environment.^1-5^ However, 30% of women in the U.S. are affected by vaginal dysbiosis, which is characterized as a polymicrobial environment colonized by pathogenic species including *Gardnerella vaginalis* and *Mobiluncus mulieris*.^6,7^ Vaginal dysbiosis contributes to an increased risk for sexually transmitted infections, pelvic inflammatory disease, and preterm birth, as well as adverse neonatal outcomes.^8-13^

Dysbiosis-associated microbes produce enzymes that degrade the cervicovaginal mucus barrier, reduce epithelial barrier integrity, and cause inflammation in local tissues.^12,14,15^ Hypotheses surrounding microbial communication to upper levels of the female reproductive tract suggest that vaginal microbes can ascend into the uterine environment, causing infection and inflammation which contributes to adverse women’s health outcomes.^16^ However, our previous work demonstrated that the pore size of cervicovaginal mucus is too small for microbes to efficiently penetrate and reach the uterus.^17^ Cervicovaginal mucus is a complex mixture of glycoproteins, ions, lipids, cells, and bacteria that protects the female reproductive tract from infections.^18-20^ Mucin proteins are crosslinked to form a heterogeneous pore structure that sterically hinders large pathogens, while the electrostatic and hydrophobic regions on these proteins adhesively interact with charged virions and particles.^18^ Given these barrier properties, we hypothesized that whole microbes would be too large to move freely through the mucus structure, preventing direct communication between bacterial cells and host cells. Thus, we sought to explore modes of microbial signaling that may directly alter tissue function in upper levels of the female reproductive tract. Specifically, we hypothesize that vaginal microbe-derived extracellular vesicles play a critical role in microbe-host communication to female reproductive tract tissues.

Bacteria-derived extracellular vesicles (bEVs) are microbe-derived, nano-sized, membrane-bound particles that enable microbe-host and microbe-microbe communication throughout the body.^21,22^ bEVs are spontaneously produced by both Gram-positive and Gram-negative bacteria, and facilitate horizontal gene transfer, defense against the host immune system, and transport of virulence factors.^21,23^ Emerging work describes the role of bEVs in mediating tissue function and disease outcomes in the lower female reproductive tract.^24-27^ These reports lay a foundation for understanding the role of bEVs in mediating gynecologic and obstetric outcomes. However, further work is necessary to understand the fate of bEVs in biological barriers (namely, cervicovaginal mucus), and how bEVs regulate tissue function in the female reproductive tract.

Here, we seek to test the hypothesis that bEVs are capable of mediating microbial communication to upper levels of the female reproductive. As such, we: (1) evaluate the mobility of both whole bacteria and bEVs through cervicovaginal mucus to assess the potential for ascension into the uterine environment; (2) assess bEV uptake by female reproductive tract cell types; and (3) establish the role of bEVs in altering tissue function relevant to the female reproductive tract. To our knowledge, our work is the first to quantify the mobility of bEVs in human cervicovaginal mucus, with direct comparisons to whole bacteria. We report changes to vaginal epithelial, endometrial, and placental cells mediated by bEVs derived from both healthy-associated and dysbiotic-associated species of human vaginal bacteria (*L. crispatus*-, *L. iners*-, *G. vaginalis*-, and *M. mulieris*). Taken together, our work points towards potential mechanisms by which vaginal bacteria-derived bEVs contribute to female reproductive tract disease, and lays a foundation for future work to develop next generation therapies for preventing and treating adverse gynecologic and obstetric outcomes.

## Materials and Methods

### Materials

Human-derived strains of *Lactobacillus crispatus* (Brygoo and Aladame, 33820), *Lactobacillus iners* (BAA-3226), *Gardnerella vaginalis* (Gardner and Dukes, 14018), and *Mobiluncus mulieris* (43064), as well as VK2/E6E7 (CRL-2616), and Kaighn’s Modification of Ham’s F-12 media, were sourced from the American Type Culture Collection (ATCC, Manassas, VA). BeWo-b30 (CCL-98) cells were received as a generous gift Dr. John Fisher at the University of Maryland, College Park, and were originally sourced from ATCC (Manassas, VA). New York City III (NYCIII) media components (HEPES, proteose peptone, sodium chloride, dextrose, and yeast extract), PKH26 red fluorescent cell linker mini kit, bovine serum albumin (BSA), Gram staining materials (crystal violet, safranin, decolorizer, and iodine), calcium chloride, poly-L-lysine, Paraformaldehyde, Amicon Ultra Centrifugal filters (10 kDa MWCO), and the Ishikawa cell line (99040201) were obtained from MilliporeSigma (Burlington, MA). Gibco™ horse serum, bicinchoninic acid assays (BCA), Amicon™ Ultra-15 Centrifugal Filter Units, Greiner Bio-One CELLSTAR µClear™ 96-well, Cell Culture-Treated, Flat-bottom Microplates, heat-inactivated fetal bovine serum (FBS), CellMask Deep Red, 4′, 6-diamidino-2-phenylindole (DAPI), keratinocyte-serum free medium, epidermal growth factor, pituitary extract, L-glutamine, minimal essential medium (MEM), and non-essential amino acids (NEAA) were ordered from ThermoFisher Scientific (Waltham, MA). V-Plex Proinflammatory Panel 1 (human) kits and Chemokine Panel 1 Gen. B kits were purchased from MesoScale Discovery (Rockville, MD). SoftDisc menstrual discs (Formerly SoftCup) were obtained from Amazon (National Landing, VA). Wiretrol^®^ disposable micropipettes were obtained from Drummond Scientific Co. (Broomall, PA). Penicillin/Streptomycin was sourced from Corning (Corning, NY). µ-Slide 18 well glass bottom imaging wells were purchased from Ibidi (Fitchburg, Wisconsin). Cell Counting Kit-8 was purchased from APExBio (Houston, TX). Formvar/Carbon 200 Mesh grids was sourced from Electron Microscopy Sciences (Hatfield, PA). 32 mL Open-Top Thickwall Polycarbonate Tubes (25 × 89mm), 38.5 mL Open-Top Thinwall Ultra-Clear Tubes (25 × 89 mm), and 5 mL Open-Top Thinwall Polypropylene Tubes (13 × 51 mm) were sourced from Beckman Coulter (Indianapolis, IN). Uranyl acetate was provided by the Laboratory for Biological Ultrastructure at the University of Maryland.

### Bacteria Culture

*L. crispatus, L. iners, G. vaginalis* and *M. mulieris* strains were cultured in particle-depleted NYCIII media. NYCIII media (0.4% w/v HEPES, 1.5% w/v proteose peptone, 0.5% w/v sodium chloride, 0.5% w/v dextrose, 2.6% w/v yeast extract) was supplemented with 10% v/v horse serum and prepared according to ATCC instructions. Particle-depleted media was prepared via ultracentrifugation using Thickwall ultracentrifuge tubes and a SW32Ti rotor (Beckman Coulter) at 100,000 x g for 12-16 h followed by 0.2 µm sterile filtration. All cultures were grown in anaerobic conditions (37°C with 5% H_2_, 10% CO_2_, 85% N_2_). Briefly, frozen stocks were plated on 1.5 % w/v agar media using the four-quadrant streak method (Day 0), then transferred to 5 mL liquid cultures using an inoculating loop (Day 3). Five mL cultures were normalized via optical density at 600 nm (OD600) prior to transfer to 50 mL cultures (Day 6). To measure OD600, 100 µL culture aliquots were taken from each culture and added to individual wells of a 96-well plate. Absorbance was read at 600 nm using a TECAN Spark® plate reader. Samples were diluted as needed to ensure an OD600 of <1. Optical densities were normalized and used to seed 50 mL cultures. Three days after seeding (Day 9), 10 µL of culture was plated on NYCIII agar plates at 10^−3^ to 10^−8^ dilutions to determine the number of colony forming units (CFUs) present in the sample. Plates were grown under anaerobic conditions for 2 days. Plates with 20-200 CFUs were considered within countable range.

### bEV Isolation

bEVs were isolated from conditioned media three days after seeding 50 mL cultures (Day 9), as previously described with modifications ^28-30^. Briefly, cultures were centrifuged at 4,000 x g for 20 mins to remove whole bacteria. The resulting supernatant was then sequentially filtered through 0.45 and 0.2 µm syringe filters. Filtered supernatant was ultracentrifuged using Thinwall ultracentrifuge tubes and a SW32Ti rotor (Beckman Coulter) at 16,000 x g for 40 mins to remove cell debris and aggregates. Supernatant was transferred to a new ultracentrifuge tube and centrifuged at 129,000 x g for 1.5 h to isolate bEVs. The bEV pellet was resuspended in PBS and centrifuged at 129,000 x g for 1.5 h to wash. Supernatant was discarded and the pellet was resuspended in the ∼1 mL of PBS. Samples were characterized immediately following isolation.

### bEV Characterization

The size, ζ-potential, and concentration of each bEV sample were measured using nanoparticle tracking analysis (NTA, ZetaView, Particle Metrix, Meerbusch) (n = 6 for each species). bEV samples were diluted 1:1000. 11 positions were run to fully characterize the sample. Parameters were set to a minimum brightness of 65, a sensitivity of 85, a frame rate of 30, and a trace length of 15. The concentration of bEVs was normalized to the initial volume of conditioned media to account for differences in the volume of PBS used for resuspension. Concentrations are reported as particles per volume of conditioned media. Samples were normalized to CFUs by dividing the particles per volume of conditioned media by CFUs per volume of conditioned media. Protein content was measured via BCA and normalized to protein per 10^8^ bEVs. Transmission electron microscopy (TEM) was completed at the University of Maryland Laboratory for Biological Ultrastructure. Samples were prepared at room temperature by depositing isolated bEVs on formvar/carbon 200 mesh grids. Samples were stained using 1% uranyl acetate. Images were taken using a HT7700 transmission electron microscope (Hitachi, Japan).

### Labeling Bacteria and bEVs

Whole bacteria were labeled using PKH26 Red Fluorescent Cell Linker Mini Kit, according to manufacturer’s instructions, with modifications as previously described.^31^ Briefly, cultures were resuspended to an OD600 of 1.3. Samples were pelleted at 4,000 x g for 20 min, and then resuspended in 2 mL of Diluent C. 4 µg of PHK26 dye was added to the mixture and then incubated for 8 min at room temperature with gentle shaking. The reaction was quenched with 2 mL of FBS. Bacteria were pelleted at 4,000 x g for 20 min and washed with PBS 2x before the pellet was resuspended in 100 µL PBS. Labeled bacteria were stored at 4°C for up to 1 month.

bEVs were labeled, as previously described.^30,32^ Equal volumes of bEVs and PKH26 reagent were mixed at a ratio of 1×10^10^ bEVs per 0.8 µg dye. The mixture was incubated for 5 min at room temperature with mixing by gentle pipetting. The reaction was quenched with an equal volume of BSA at a ratio of 1 mg per 1×10^10^ bEVs. Samples were immediately placed on top of a 2 mL 20% sucrose cushion and pelleted at 100,000 x g for 2 h in Thinwall ultracentrifuge tubes and a SW55Ti rotor (Beckman Coulter). Pellets were resuspended in 5 mL of PBS and washed in a 15 mL 10 kDa MCWO filter. Samples were analyzed via NTA to determine final concentration.^30^ Labeled bEVs were stored at 4°C for up to 1 week until use.

### Cervicovaginal Mucus Collection

Human cervicovaginal mucus (CVM) was collected in accordance with protocol #2043110-3, as approved by the University of Maryland Institutional Review Board. Participants were determined to be in the luteal phase based on reported dates of menstruation and urine ovulation tests. Participants self-collected CVM using SoftDisc menstrual devices, as previously described.^17,33^ Briefly, participants inserted the SoftDisc into the vagina for up to 1 min, and then removed the disc in a twisting motion to collect CVM. The SoftDisc was then placed into a 50 mL conical and spun at 300 x g for 5 min to collect CVM. Samples were characterized via wet mount, Nugent score, and pH (**Supp. Table 1**). For wet mounts, <10 µL of CVM was placed on a slide and smoothed using a wiretrol. 10 µL normal saline was pipetted onto the sample to seal a coverslip. Slides were imaged on a ZEISS Axiovert 5 equipped with a 100x objective. Nugent scores were assigned based on Gram staining CVM smears.^17^ The pH of each mucus sample was measured using a micro pH probe (MI-4146b, MicroElectrodes, Inc).

### Multiple-Particle Tracking

Multiple-particle tracking (MPT) was conducted to assess the mobility of whole bacteria and bEVs in human CVM, as previously described.^17,30^ Three-dimensional wells were constructed using a 6 mm diameter hole punch through two layers of electrical tape adhered to a glass slide. 20 μL of fresh CVM was added to the well along with 1.5 μL of fluorescently labeled bacteria or bEVs. Particle mobility was recorded for 10-15 s at room temperature on a ZEISS Axiovert 5 equipped with a 100x objective. An Axiocam 305 Color camera was used to record videos at a frame rate of 66.6 frame/s. For each CVM sample, a minimum of 5 videos were taken for each particle type. The mean squared displacement (MSD) for each individual particle was calculated using image processing software in MATLAB, as previously described (n = 10 CVM samples).^17,30^

### Cell Culture

VK2/E6E7 vaginal epithelial cells were cultured in keratinocyte-serum free medium supplemented with 0.1 ng/ml human recombinant EGF, 0.05 mg/ml bovine pituitary extract, and additional 44.1 mg/L calcium chloride, according to manufacturer’s instructions. Experiments utilized passages 5-18. Ishikawa endometrial cells were cultured in MEM media supplemented with 2 mM L-glutamine, 1% v/v NEAA, and 5% v/v FBS, according to manufacturer’s instructions. Experiments utilized passages 6-10. For VK2/E6E7 and Ishikawa cell lines, media was exchanged every 48-72 h until confluent. BeWo-b30 placental cells were cultured in F-12K supplemented with 10% v/v FBS and 1% v/v penicillin/streptomycin. Experiments utilized passages 24-33. Media was exchanged every 24-48 h until confluency. All cell lines were maintained at 37°C, 5% CO_2_. Cells were passaged at 85-95% confluency.

### bEV Uptake

Cells were seeded at 0.04 × 10^6^ cells per well in a black walled 96-well plate and allowed to adhere overnight. Media was exchanged immediately before dosing wells with 2500 labeled bEVs/cell. Uptake was measured 2, 8, 12, and 24 h after dosing (n = 8 wells/timepoint). Cells were washed with PBS 3x before incubation with 1% Triton for 1 h, as previously described.^34^ Fluorescent readings were taken at 530:567 nm to determine internalized bEV concentrations. Standard curves were created using known concentrations of bEVs in 1% Triton. Values below the limit of detection were assigned a value of zero.

For confocal imaging, µ-Slide glass bottom 18-well plates were treated with poly-L-lysine for 20 mins and dried overnight. Cells were seeded at 0.04 × 10^6^ cells/well and allowed to adhere overnight. As above, media was exchanged immediately prior to dosing with 2500 labeled bEVs/cell. After 24 h, cells were labeled with CellMask Deep Red membrane stain for 5-10 mins, according to manufacturer’s instructions. Cells were then fixed via incubation with 2% w/v formaldehyde at 37°C for 5 mins and washed with PBS 3x. Cells were incubated in 300 nM DAPI for 5 mins and washed with PBS according to manufacturer’s instructions. Cells were imaged using a LSM 980 Laser Scanning Confocal microscope (Zeiss, Oberkochen, Germany; n = 3 images per well, 3 wells per condition).

### Cell Viability

Cells were seeded at 0.04 × 10^6^ cells per well in a 96-well plate and allowed to adhere overnight. Media was exchanged immediately before dosing wells with 50, 500, or 5000 bEVs per cell. After 24 h, Cell Counting Kit-8 (CCK-8) was used according to manufacturer’s instructions. Briefly, 10 μL of assay reagent was added to each well. Samples were incubated at 37°C for 1 h. Viability was determined by measuring the OD450 and normalizing to background (OD600). Cell viability was normalized to vehicle control (PBS) (n = 8 per treatment group).

### Multiplex Assays

Cells were seeded at 0.24 × 10^6^ in a 24-well plate and allowed to adhere overnight. Media was exchanged immediately before dosing wells with 5000 bEVs per cell. Samples were incubated 24h, after which supernatants were collected and stored at -80°C until use. IL-1β, IL-6, IL-8, IL-10, TNF-α, MIP-1β, IP-10, and MIP-1α cytokines were measured using custom V-PLEX plates, according to the manufacturer’s instructions. Samples were diluted as needed in the supplied sample buffer, and 50 µL of diluted sample was added to appropriate wells. Plates were incubated at RT for 2h while shaking at 700 rpm. After washing 3x, 25 µL of detection antibody solution was added to each well, and the plate was incubated at RT while shaking for 2h at 700 rpm. The plate was washed 3x and 150 µL of read buffer was added to each well by reverse pipetting. The plate was immediately read using the MSD Meso Quickplex SQ 120MM Imager. Cytokine concentration (n=4-8 per condition) was calculated based on a standard curve run on each plate using Discovery Workbench software from MSD.

### Statistical Analysis

GraphPad Prism was used for statistical analysis. For bEV characterizations, and multiplex experiments, 1-way analysis of variance (ANOVA) testing with multiple comparisons was used with significance at *p* < 0.05. Viability was analyzed via 2-way ANOVA with comparisons to vehicle control. Cellular uptake experiments were analyzed via 2-way ANOVA, with multiple comparisons between groups at each timepoint. Replicates were assigned as zero if fluorescent values were below the standard curve. For multiple-particle tracking analysis, two-tailed Mann-Whitney non-parametric tests were performed to compare whole bacteria and bEV mobility in each individual participant, as previously described.^35^ For characterization, uptake, and viability experiments, outliers were removed according to Grubb’s test with significance at *p* < 0.05. Data are presented as mean ± SEM.

## Results

*Physical characteristics of bacteria-derived extracellular vesicles differ based on cell of origin* Throughout this work, we evaluated differences in bEVs isolated from human-derived, commercially available strains of *L. crispatus, L. iners, G. vaginalis*, and *M. mulieris. L. crispatus* is used as a representative of a healthy vaginal microbiome; *L. iners* representative of an intermediate microbiome; *G. vaginalis* representative of dysbiosis; and *M. mulieris* representative of advanced dysbiosis. We first sought to examine potential differences in the physical characteristics of bEVs from these four strains. Differences in size, surface charge (ζ-potential), and production may dictate bEV interactions with the vaginal microenvironment and female reproductive tract. bEVs were confirmed to be membrane-bound particles via TEM (**Figure 1A, Supp. Figure 1**). bEV concentration was highest from *M. mulieris* cultures, followed by *L. crispatus* > *L. iners* > *G. vaginalis. G. vaginalis*-derived bEVs were significantly lower in concentration (1.30 ± 0.089 × 10^9^) compared to isolated bEVs from *L. crispatus* (2.05 ± 0.16 × 10^9^ particles/mL, *p* = 0.0156) and *M. mulieris* cultures (2.14 ± 0.24 × 10^9^ particles/mL, *p* = 0.0110) (**Figure 1B-C**). When normalized to CFUs, bEV production was highest from *L. crispatus* cultures, followed by *L. iners* > *G. vaginalis* > *M. mulieris. L. crispatus* cultures (7.15 ± 1.35 × 10^9^ bEVs/CFU) had significantly higher bEV production compared to *G. vaginalis* (1.67 ± 0.36 × 10^9^ bEVs/CFU; *p* = 0.0048) and *M. mulieris* cultures (0.24 ± 0.05 × 10^9^ bEVs/CFU, *p* = 0.0015). *L. iners* cultures (5.49 ± 1.42 × 10^9^ bEVs/CFU) had higher bEVs production compared to *M. mulieris* cultures (*p* = 0.0145) (**Figure 1D**). No differences in bEV protein content were observed (**Figure 1E**). *G. vaginalis*-derived bEVs (134.0 ± 2.293 nm) and *M. mulieris*-derived bEVs (131.7 ± 2.809 nm) were smaller than *L. crispatus*- and *L. iners*-derived bEVs (143.6 ± 1.327 nm, *p ≤* 0.0182; 154.5 ± 1.785 nm, *p* < 0.0001, respectively) (**Figure 1F**). *G. vaginalis*-derived bEVs had a more neutral ζ-potential (−26.5 ± 1.221 mV) compared to *L. crispatus*-derived bEVs (−30.85 ± 0.537 mV, *p* = 0.0317), *L. iners*-derived bEVs (−31.90 ± 1.070 mV, *p* = 0.0072), and *M. mulieris*-derived bEVs (−31.30 ± 1.107 mV, *p* = 0.0257) (**Figure 1G**).

**Figure 1:**
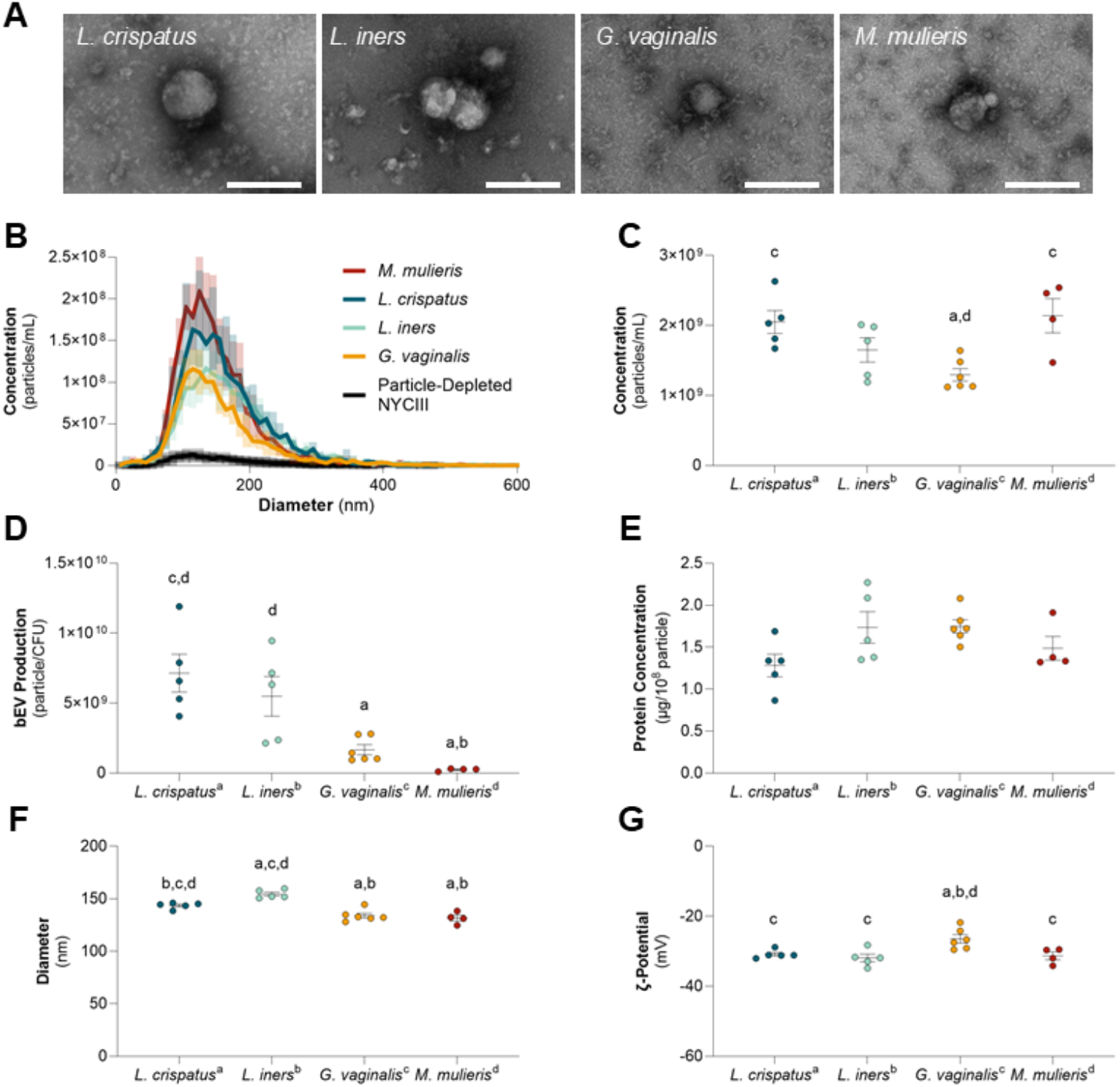
Physical characteristics of vaginal bacteria-derived bEVs. Analysis was completed via nanoparticle tracking analysis. (**A**) Isolated bEVs were confirmed to be membrane-bound particles via TEM. Scale bars denote 200 nm. (**B, C**) *G. vaginalis*-derived bEVs were significantly lower in concentration (1.30 ± 0.089 × 10^9^) compared to isolated bEVs from *L. crispatus* (2.05 ± 0.16 × 10^9^ particles/mL, *p* = 0.0156) and *M. mulieris* cultures (2.14 ± 0.24 × 10^9^ particles/mL, *p* = 0.0110). (**D**) *L. crispatus* cultures (7.15 ± 1.35 × 10^9^ bEVs/CFU) had significantly higher bEV production compared to *G. vaginalis* cultures (1.67 ± 0.36 × 10^9^ bEVs/CFU; *p* = 0.0048) and *M. mulieris* cultures (0.24 ± 0.05 × 10^9^ bEVs/CFU, *p* = 0.0015). *L. iners* cultures (5.49 ± 1.42 × 10^9^ bEVs/CFU) had higher bEVs production compared to *M. mulieris* cultures (*p* = 0.0145). (**E**) No differences in bEV protein content were observed. (**F**) *G. vaginalis* bEVs (134.0 ± 2.293 nm) and *M. mulieris*-derived bEVs (131.7 ± 2.809 nm) were smaller than *L. crispatus*- and *L. iners*-derived bEVs (143.6 ± 1.327 nm, *p ≤* 0.0182; 154.5 ± 1.785 nm, *p* < 0.0001, respectively). (**G**) *G. vaginalis*-derived bEVs had a more neutral ζ-potential (−26.5 ± 1.221 mV) compared to *L. crispatus*-derived bEVs (−30.85 ± 0.537 mV, *p* = 0.0317), *L. iners*-derived bEVs (−31.90 ± 1.070 mV, *p* = 0.0072), and *M. mulieris*-derived bEVs (−31.30 ± 1.107 mV, *p* = 0.0257). Values are reported as mean ± SEM. Statistical significance (*p* ≤ 0.05) is represented by a letter corresponding to the species of comparison (a, *L. crispatus;* b, *L. iners*; c, *G. vaginalis*; d, *M. mulieris*). (n = 6 per group, 2 outliers removed from the *M. mulieris* group in accordance with Grubb’s outlier test due to size and concentration).

### bEVs show increased mobility in human cervicovaginal mucus compared to whole bacteria

To test the hypothesis that bEVs can penetrate human CVM more efficiently than whole bacteria, we used multiple-particle tracking technology to determine individual particle mobility through CVM. To ensure comprehensive sampling, an equal number of participants samples with high and low pH (pH above or below 4.2) CVM were utilized (**Supp. Table 1**). Nugent scores for each sample (n = 10) were assigned based on Gram-stained mucus smears (**Supp. Table 1**). Across all samples, we observed increased mobility by bEVs, as compared to whole bacteria. Geometric means of the mean squared displacement were determined using the 1 second time point (**Figure 2A**). For *L. crispatus* whole bacteria and bEVs, all CVM samples showed limited mobility of whole bacteria compared to bEVs (*p* ≤ 0.0076) (**Figure 2B**). For *L. iners*, 9/10 participants showed limited mobility of whole bacteria compared to bEVs (*p* ≤ 0.0001). One CVM sample demonstrated trapping of both bEV and whole microbes (**Figure 2C**, Sample 7, *p* = 0.0621). For *G. vaginalis*, 6 out of 10 samples showed increased mobility of bEVs over whole bacteria (*p* ≤ 0.01). 2 out of 10 samples (Samples 7, 10) showed decreased mobility of both whole microbes and bEVs. For *M. mulieris*, 9 out of 10 samples showed limited mobility of whole bacteria compared to bEVs (*p* ≤ 0.0001, **Figure 2E**). Given the increased mobility of bEVs in CVM, compared to whole bacteria, we hypothesize that bEVs represent an important mediator of microbe-host communication in the female reproductive tract. We next sought to evaluate the interactions of bEVs with female reproductive tract tissues, moving from vaginal epithelial cells, to endometrial cells, and, finally, to the placenta.

**Figure 2:**
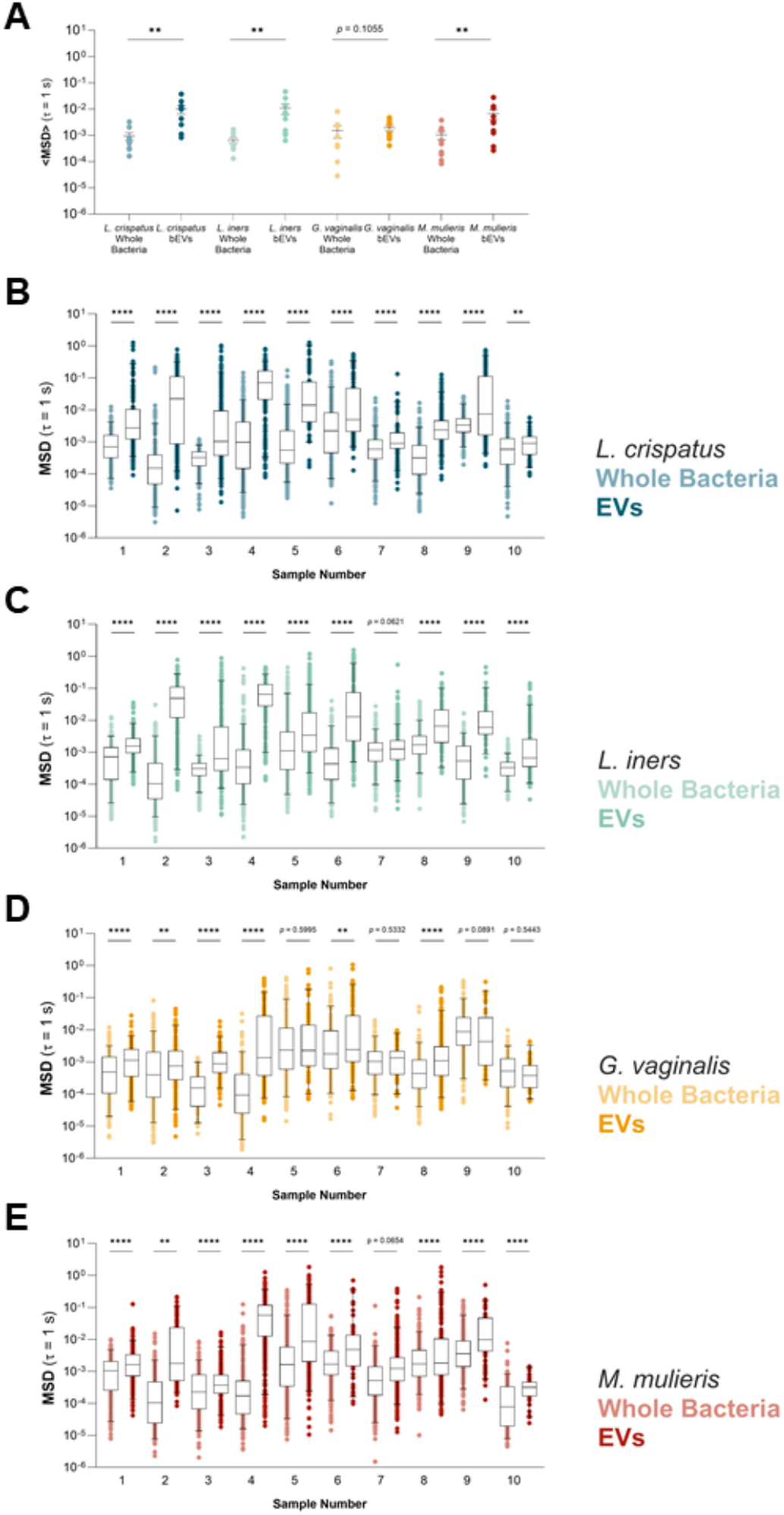
Whole bacteria show limited mobility across all species. (**A**) Geometric means of the average mean squared displacement at 1 second for all samples. bEVs demonstrate increased mobility compared to whole bacteria (**B**) All CVM samples showed limited mobility of *L. crispatus* whole bacteria, compared to bEVs (*p* ≤ 0.01). (**C**) 9/10 samples showed limited mobility of *L. iners* whole bacteria compared to bEVs (*p* ≤ 0.0001). (**D**) 6/10 samples showed increased mobility of *G. vaginalis* bEVs compared to whole bacteria (*p* ≤ 0.01). (**E**) 9/10 samples demonstrated limited mobility of whole bacteria compared to bEVs (p ≤ 0.0001). Box and Whiskers are shown as 5-95% confidence intervals. Two-tailed Mann-Whitney non-parametric tests were performed to compare whole bacteria and bEV mobility in each individual participant. n = 10 samples.

### Vaginal epithelial cells exhibit a pro-inflammatory response after treatment with G. vaginalis- and M. mulieris-derived bEVs

Based on the evidence that bEVs diffuse through CVM, we sought to determine vaginal epithelial response due to bEVs. After 24 h incubation, confocal microscopy verified bEV presence within cytosol (**Figure 3A, Supp. Figure 2**). Uptake assays determined that there were significant differences in the uptake based on bEV type (**Figure 3B**). Specifically, vaginal epithelial cells internalized *M. mulieris*-derived bEVs (23.88 ± 2.43%) significantly more than *L. crispatus*-derived bEVs (13.30 ± 1.99%, *p* = 0.0001), *L. iners*-derived bEVs (15.52 ± 2.13%, *p* = 0.0074), or *G. vaginalis*-derived bEVs (13.86 ± 2.03%, *p* = 0.0005) at 24 h (**Figure 3C**).

**Figure 3.**
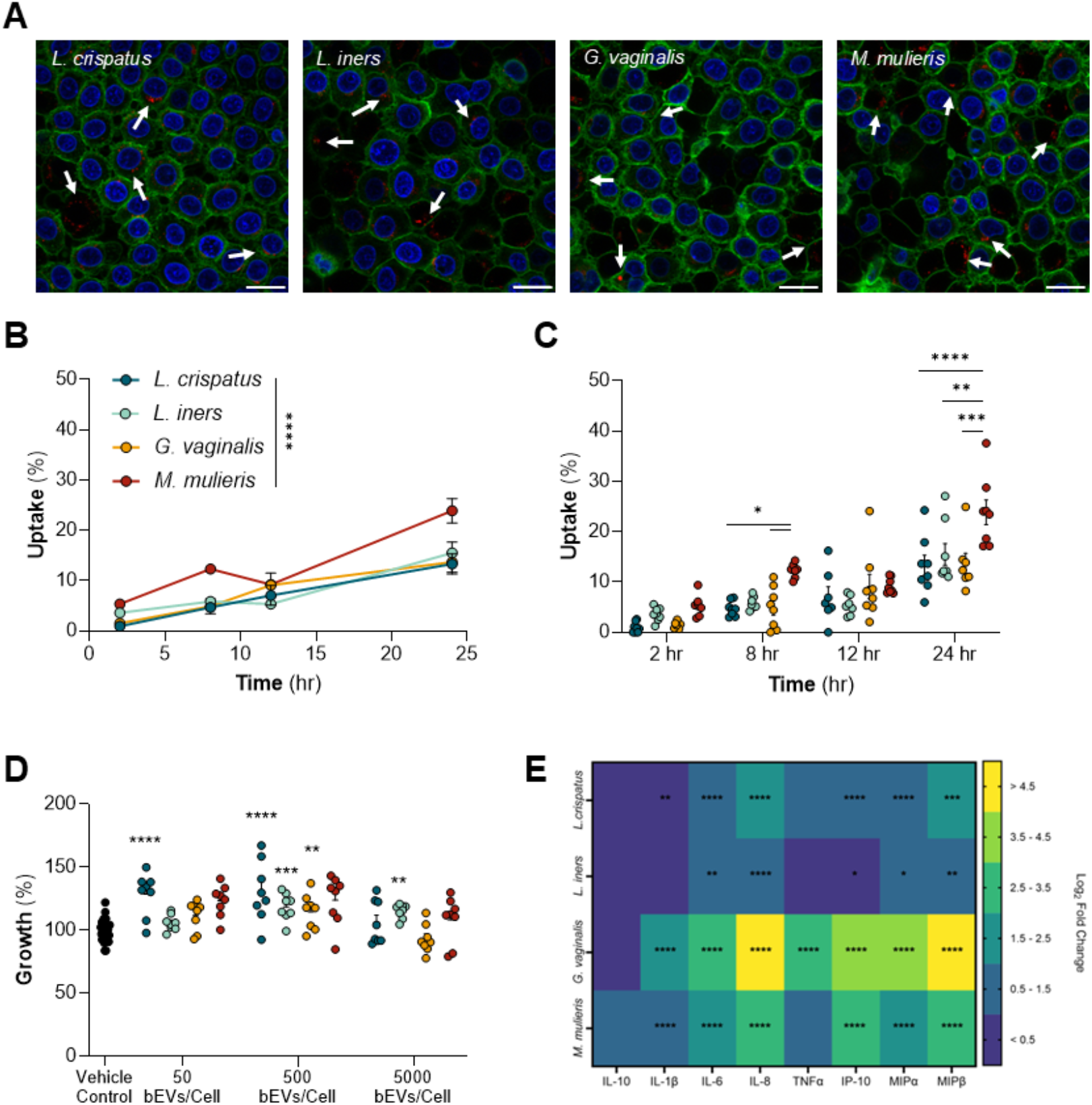
Vaginal epithelial cells exhibit a pro-inflammatory response after treatment with G. vaginalis- and M. mulieris-derived bEVs. (**A**) Confocal images reveal uptake of bEVs to the cytosol. Scale bars denote 20 µm. (**B**) Based on 2-way ANOVAs, uptake assays determined that there were significant differences in the uptake based on bEV type. (**C**) Vaginal epithelial cells internalized *M. mulieris*-derived bEVs (23.88 ± 2.43%) significantly more than *L. crispatus*-derived bEVs (13.30 ± 1.99%, *p* = 0.0001), *L. iners*-derived bEVs (15.52 ± 2.13%, *p* = 0.0074), or *G. vaginalis*-derived bEVs (13.86 ± 2.03%, *p* = 0.0005) at 24 h (n=8 per dosage per timepoint. Replicates below the standard curve were assumed to be 0%.) (**D**) Vaginal epithelial cells did not exhibit a decrease in viability after 24 h incubation with bEVs of any dosage or species. (n=8 for treatment group, n=16 for vehicle control). (**E**) VK2/E6E7 cells exhibit a pro-inflammatory response after treatment with G. vaginalis- and M. mulieris-derived bEVs (n=8). Statistics for uptake and viability were performed using 2-way ANOVA. Statistics for cytokine production were performed using 1-way ANOVA for each cytokine using raw cytokine concentrations.

Although we observed no cytotoxicity in vaginal epithelial cells after 24 h exposure to bEVs from any species at any concentration (**Figure 3D**), we sought to evaluate the change in cell function by measuring cytokine and chemokine production after bEV exposure. *G. vaginalis*-derived bEVs resulted in increased log_2_ fold change in IL-1β (1.63 ± 0.06, *p* < 0.0001), IL-6 (2.68 ± 0.18, *p* < 0.0001), IL-8 (5.53 ± 0.09, *p* < 0.0001), TNFα (3.05 ± 0.15, *p* < 0.0001), IP-10 (4.47 ± 0.07, *p* < 0.0001), MIPα (3.67 ± 0.08, *p* < 0.0001), and MIPβ (5.25 ± 0.14, *p* < 0.0001), compared to the vehicle control. *M. mulieris*-derived bEVs resulted in increased IL-1β (0.85 ± 0.05, *p* = 0.0425), IL-6 (1.93 ± 0.18, *p* = 0.0001), IL-8 (3.36 ± 0.12, *p* = 0.0004), IP-10 (2.82 ± 0.10, *p* < 0.0001), and MIPα (2.38 ± 0.16, *p* < 0.0001), compared to the vehicle control. *L. crispatus*-derived bEVs resulted in a slight increase in IL-6 (1.41 ± 0.11, *p* = 0.0091) and MIPα (1.48 ± 0.17, *p* = 0.0294). No changes in cytokines were seen in response to treatment with *L. iners*-derived bEVs (**Figure 3E, Supp. Figure 4**). These data suggest species-specific responses based on bEV signaling.

### bEVs result in a differential expression of IL-8 in endometrial cells

Moving from the vaginal environment into the uterine environment, bEVs will encounter endometrial tissue. Thus, we investigated the endometrial cell response to vaginal bacteria-derived bEVs. Microscopy verified bEV presence within cytosol at 24 h (**Figure 4A, Supp. Figure 4**). Again, uptake assays determined that there were significant differences in the uptake based on bEV type (**Figure 4B**). *L. crispatus*-derived bEVs (14.06 ± 0.91%) exhibited the highest uptake at 24 h compared to *L. iners* bEVs (9.97 ± 0.67%, *p* < 0.0001), *G. vaginalis*-derived bEVs (1.468 ± 0.52%, *p* < 0.0001), and *M. mulieris*-derived bEVs (7.09 ± 0.73%, *p* < 0.0001) (**Figure 4C**). bEVs did not lead to cytotoxicity in endometrial cells at any of the tested doses (**Figure 4D**).

When investigating cytokine and chemokine production, treatment with *G. vaginalis*-derived bEVs resulted in increased IL-8 (1.27 ± 0.10, *p* < 0.0001) compared to the vehicle control. No changes in cytokines were seen in response to treatment with *L. crispatus-, L. iners*-, or *M. mulieris*-derived bEVs (**Figure 4E, Supp. Figure 5**).

**Figure 4.**
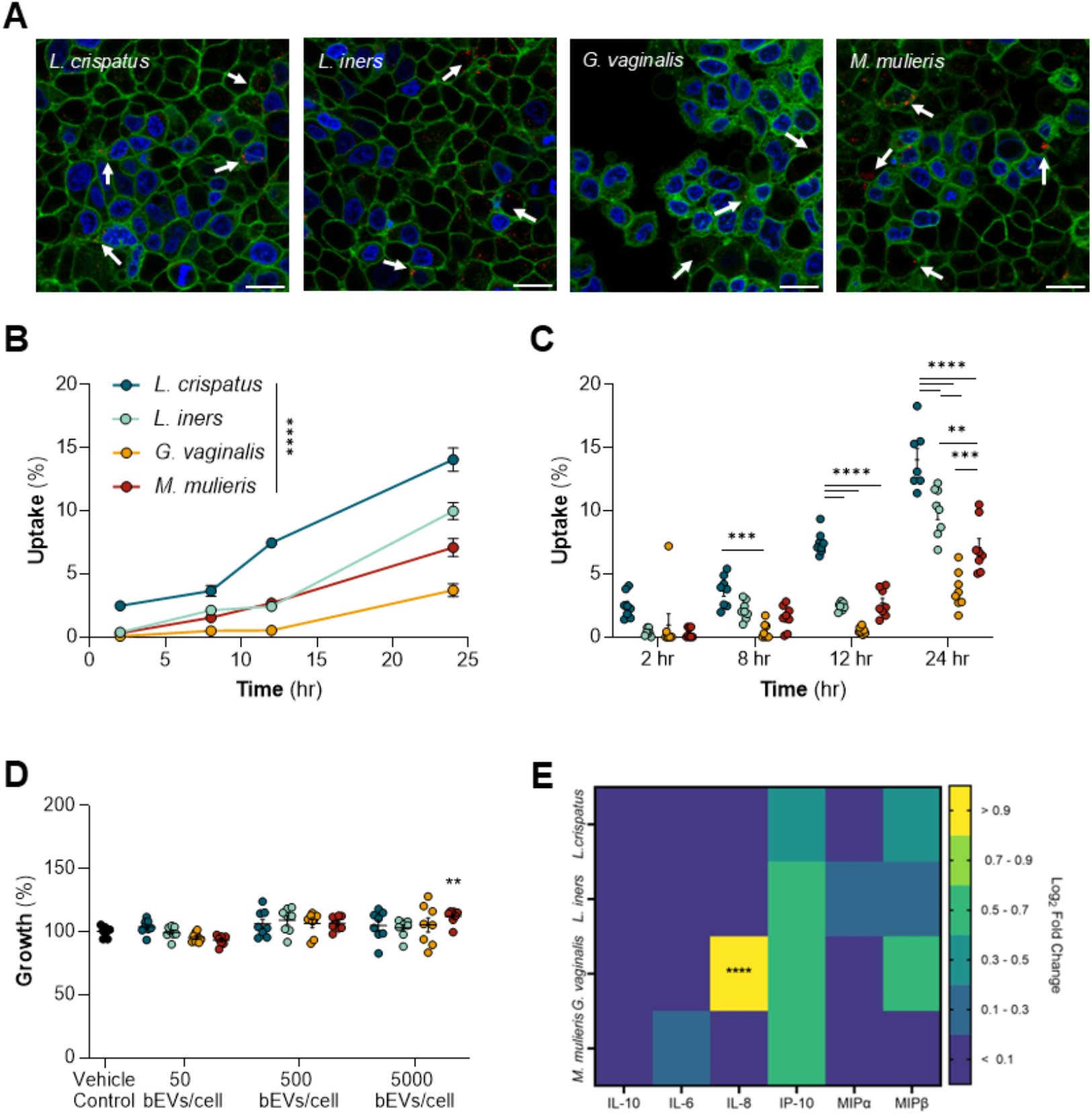
Endometrial cells exhibit a change in IL-8 production after treatment with G. vaginalis and M. mulieris. (**A**) Confocal images reveal uptake of bEVs to the cytosol. Scale bars denote 20 µm. (**B**) Based on 2-way ANOVAs, uptake assays determined that there were significant differences in the uptake based on bEV type. (**C**) L. crispatus-derived bEVs (14.06 ± 0.91%) saw the highest uptake at 24 h compared to L. iners-derived bEVs (9.97 ± 0.67%, p < 0.0001), G. vaginalis-derived bEVs (1.468 ± 0.52%, p < 0.0001), and M. mulieris-derived bEVs (7.09 ± 0.73%, p < 0.0001) (n=8 per dosage per timepoint. Replicates below the standard curve were assumed to be 0%.) (**D**) Endometrial cells did not exhibit a decrease in viability after 24 h incubation with bEVs of any dosage or species. (n=8) (**E**) Endometrial cells exhibit changes in IL-8 production after dosage with G. vaginalis- and M. mulieris-derived bEVs (n=9). Treatment with G. vaginalis-derived bEVs resulted in an increase in log_2_ fold change (1.27 ± 0.10, p < 0.0001). Statistics for uptake and viability were performed using 2-way ANOVA. Statistics for cytokine production were performed using 1-way ANOVA for each cytokine using the raw concentrations.

*G. vaginalis-derived bEVs increase pro-inflammatory cytokine production in placental cells* Given the critical role of the vaginal microbiome in pregnancy and neonatal outcomes, we next investigated bEV interactions with the placenta. Uptake experiments verified bEV presence within cytosol at 24 h (**Figure 5A, Supp. Figure 6**), and uptake assays determined significant differences in the uptake based on bEV type (**Figure 5B**). *L. crispatus*-derived bEVs were most internalized after 24 h (41.58 ± 2.502%) (**Figure 5C**). *G. vaginalis*-derived bEVs exhibited the second highest uptake, reaching 22.21 ± 0.988%. *L. iners-* and *M. mulieris*-derived bEVs saw lower uptake, at 13.890 ± 1.244% and 5.097 ± 0.544% respectively (**Figure 5C**). Finally, we sought to determine placental cell viability and inflammatory cytokine secretion in response to bEVs. We did not observe placental cell cytotoxicity caused by any of the tested bEV doses (**Figure 5D**).

**Figure 5.**
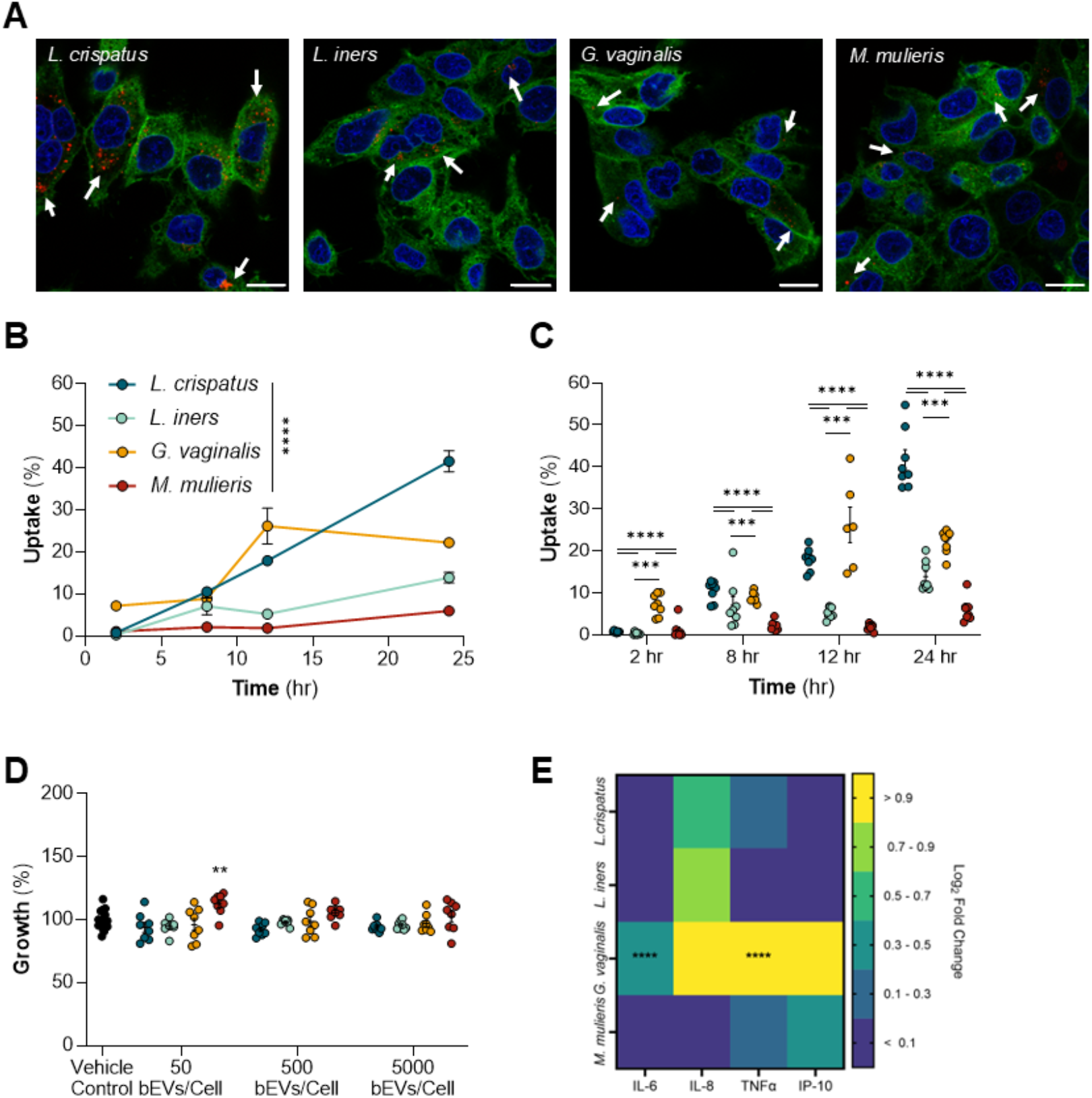
Placental cells exhibit an increase in proinflammatory cytokine production after treatment with *G. vaginalis*-derived bEVs. (**A**) Confocal images show uptake of bEVs to the cytosol. Scale bars denote 20 µm. (**B**) Based on 2-way ANOVAs, uptake assays determined that there were significant differences in the uptake based on bEV type. (**C**) *L. crispatus*-derived bEVs were most internalized after 24 h (41.58 ± 2.502%). *G. vaginalis*-derived bEVs exhibited the second highest uptake, reaching 22.21 ± 0.988%. *L. iners*- and *M. mulieris*-derived bEVs saw lower uptake, at 13.890 ± 1.244% and 5.097 ± 0.544% respectively (n=8 per dosage per timepoint. Replicates below the standard curve were assumed to be 0%.) (**D**) BeWo-b30 cells did decrease in viability after 24 h incubation with bEVs of any dosage or species. (n=8 for treatment group, n=16 for vehicle control). (**E**) Multiplex assays studies revealed significant increases of the log_2_ fold change in IL-6 (0.43 ± 0.06, *p* = 0.0010) and TNFα (1.91 ± 0.014, *p* < 0.0001) after dosage with *G. vaginalis*-derived bEVs. Statistics for uptake and viability were performed using 2-way ANOVA. Statistics were performed using 1-way ANOVA for raw concentrations of each cytokine.

When investigating cytokine and chemokine production, treatment with *G. vaginalis*-derived bEVs resulted in increased IL-6 (0.43 ± 0.06, *p* = 0.0010) and TNFα (1.91 ± 0.014, *p* < 0.0001) compared to the vehicle control. No changes in cytokines were seen in response to treatment with *L. crispatus-, L. iners*-, or *M*. mulieris-derived bEVs (**Figure 5E, Supp. Figure 7**).

## Discussion

The vaginal microbiome is significantly implicated in female reproductive health outcomes.^2,13,36^ Extensive clinical work establishes that dominance by *Lactobacillus* spp. in the vaginal microbiome is associated with healthy outcomes, whereas a polymicrobial environment leads to increased risk for gynecologic, obstetric, and neonatal disease.^36^ While the proximity of vaginal bacteria to the vaginal epithelium and cervix may facilitate direct microbial signaling, how the vaginal microbiome affects upper levels of the female reproductive tract is less well understood. One hypothesis surrounding microbial signaling to the upper female reproductive tract is the ascension of vaginal bacteria into the uterine environment, which could lead to functional changes to uterine and placental tissues. However, prior work from our group suggests that the CVM barrier would not permit the ascension of whole bacteria from the vagina into the uterus.^17^ Instead, we hypothesize that communication between the vaginal microbiome and the female reproductive tract is primarily modulated by bEVs derived from the vaginal microbiota.

Here, we isolated bEVs from human-derived, commercially available strains of *L. crispatus, L. iners, G. vaginalis*, and *M. mulieris*. Consistent with previous work, we observe the diameter of bEVs to be 50-250 nm.^25,27,37,38^ We observed differences in bEV production rates, with *L. crispatus* and *L. iners* demonstrating higher productivity (bEVs/CFU) compared to dysbiotic species (*G. vaginalis* and *M. mulieris*). In the transition from a healthy to a dysbiotic vaginal environment, this loss of commensal or probiotic bEVs may reduce positive communication, increasing risk for adverse reproductive outcomes. Understanding differences in bEV activity within the vaginal microenvironment is critical to better understanding and treating female reproductive diseases—both in terms of promoting healthy interactions and preventing dysbiotic interactions.

In order to deliver cargoes to female reproductive tissues, bEVs must be able to move through the protective CVM barrier. Previous work demonstrates a critical relationship between mucus barrier properties and risk for gynecologic and obstetric disease. ^17-19,39-41^ However, we hypothesized that, even weakened barrier properties of dysbiotic mucus would prohibit direct interactions between bacterial cells and host cells. Indeed, we observed a significantly decreased mobility of whole microbes in comparison to bEVs of the same species, suggesting that it is unlikely whole microbes can directly communicate with upper reproductive tissues. The mobility of bEVs in CVM suggests the potential for these bacterial byproducts to reach upper reproductive tract tissues and facilitate signaling between vaginal microbiota and female reproductive tract tissues. Our results support emerging work in the EV and bEV fields demonstrating the ability of these cell-derived nanoparticles to cross biological barriers, facilitating long-distance communication in the body.^42-46^

Vaginal epithelial cells form a tissue barrier to infections in the female reproductive tract. In our work, bEVs from all species were internalized by vaginal epithelial cells. *M. mulieris*-derived bEVs were more significantly internalized compared to other microbe-derived bEVs. As a dysbiotic species, the improved uptake of *M. mulieris*-derived bEVs may be especially detrimental to reproductive tissues. Additionally, the increased production of IL-6 by vaginal epithelial cells in response to *G. vaginalis*-derived and *M. mulieris*-derived bEVs is consistent with clinical reports of vaginal dysbiosis.^47^ Our data support the hypothesis that bEVs may be a mediator between vaginal microbes and reproductive tissues, increasing risk of adverse gynecologic and obstetric outcomes.

While some work has investigated bEV-mediated changes to lower female reproductive tract cells, no prior work has investigated vaginal bEV-mediated changes to endometrial cells. The endometrium plays a critical role in implantation, requiring a balance of pro- and anti-inflammatory signals.^48^ Previous work reported the presence of bacteria within endometrial samples, revealing an association between dysbiosis and decreased implantation rates.^49,50^ Our data demonstrate the uptake of bEVs by endometrial cells, as well as functional changes to inflammatory cytokine production. In particular, *G. vaginalis*-derived bEVs significantly increased the production of IL-8 in endometrial cells.^51^ These findings suggest that bEVs may, in part, mediate endometrial function.

Beyond vaginal epithelial and endometrial cells, microbial interactions with the placenta may impact preterm birth, as well as fetal programming.^42,52-55^ Even in full-term infants, maternal vaginal dysbiosis is associated increased risk for respiratory distress, low birthweight, neonatal sepsis, and admission to the neonatal intensive care unit.^56^ bEVs have been shown to directly degrade placental barrier properties, demonstrating their ability to modulate tissue function.^42^ We report *in vitro* placental uptake of bEVs from all four species over the course of 24 h. Notably, *L*. crispatus-derived bEVs were most significantly internalized, suggesting the need for further understanding surrounding lipid and protein composition that dictate *in vivo* interactions. Functionally, in a dysbiotic environment, the loss of probiotic-derived bEVs may exacerbate pro-inflammatory responses and negatively affect reproductive tissue function. Understanding microbe-host signaling in the placenta is critical to developing strategies that promote a healthy environment and healthy long-term development in offspring. Our observed increase in TNF-α production after exposure to *G. vaginalis*-derived bEVs may have implications in cell stress and death, which has been linked to preterm birth and other placental diseases.^57^ Similarly, the reported IL-6 responses may indicate that bEVs initiate placental distress and inflammation which contributes to preterm birth.^58-60^ Recent evidence of bEVs both in human placentas and infant meconium make bEV-placenta interactions a key area of interest.^43,44^

In summary, our work is the first to examine the mobility of whole bacteria and bEVs in CVM, and is the first to explore the interactions between vaginal microbe-derived bEVs and upper female reproductive tract cells. Our data suggests a novel role for bEVs in mediating microbe-host signaling relevant to gynecologic and obstetric diseases. While whole bacteria are unable to penetrate vaginal mucus, vaginal bacteria-derived bEVs can diffuse more freely to facilitate microbe-host communication in upper levels of the female reproductive tract. We also demonstrated that cells in all levels of the female reproductive tract were able to internalize bEVs from all four vaginal microbe species. Beyond uptake, we observe that bEVs mediate inflammatory response in reproductive cells, with relevance to bacterial vaginosis, endometriosis, preterm birth, and placental programming. Future work should evaluate bEV-mediated microbe-host interactions using more complex models of the female reproductive tract, understand species- and strain-level differences in terms of bEV composition, and leverage findings to develop next generation therapies for gynecologic and obstetric indications.

## Data Availability

All data generated or analyzed during this study are included in this published article and its supplementary information file.

## Author Contributions

DS and HCZ conceptualized the study. DS, APP, and YC performed and analyzed experiments. DS wrote the original draft. DS, APP, and HCZ reviewed and edited the manuscript. All authors read and approved the final manuscript.

## Acknowledgements

This work was funded by a Faculty-Student Research Award from the Graduate School, University of Maryland (HCZ), the Minta Martin Foundation (HCZ) We acknowledge the support of the University of Maryland, Baltimore, Institute for Clinical & Translational Research (ICTR) and the National Center for Advancing Translational Sciences (NCATS) Clinical Translational Science Award (CTSA), UM1TR004926, as well as the University of Maryland Strategic Partnership: MPowering the State (MPower) (DS, HCZ). DS was supported by the Microbiome Summer Support Fellowship through the Center of Excellence in Microbiome Sciences. APP was supported by the Clark Doctoral Fellows Program through the A. James Clark School of Engineering. Purchase of the Zeiss LSM 980 Airyscan 2 was supported by Award Number 1S10OD025223-01A1 from the National Institute of Health. Funders played no role in the study design, data collection, analysis and interpretation of data, or the writing of the manuscript. Finally, we would like to thank Dr. Senta Kapnick from Dr. Christopher Jewell’s Laboratory and Dr. Chen-Yu Chen from Dr. William Bentley’s Laboratory for their insight and use of equipment throughout this work.

