## Supplementary Information for "Vaginal bacteria-derived extracellular vesicles diffuse through human cervicovaginal mucus to enable bacterial signaling to upper female reproductive tract tissues"

### Supplementary Figures

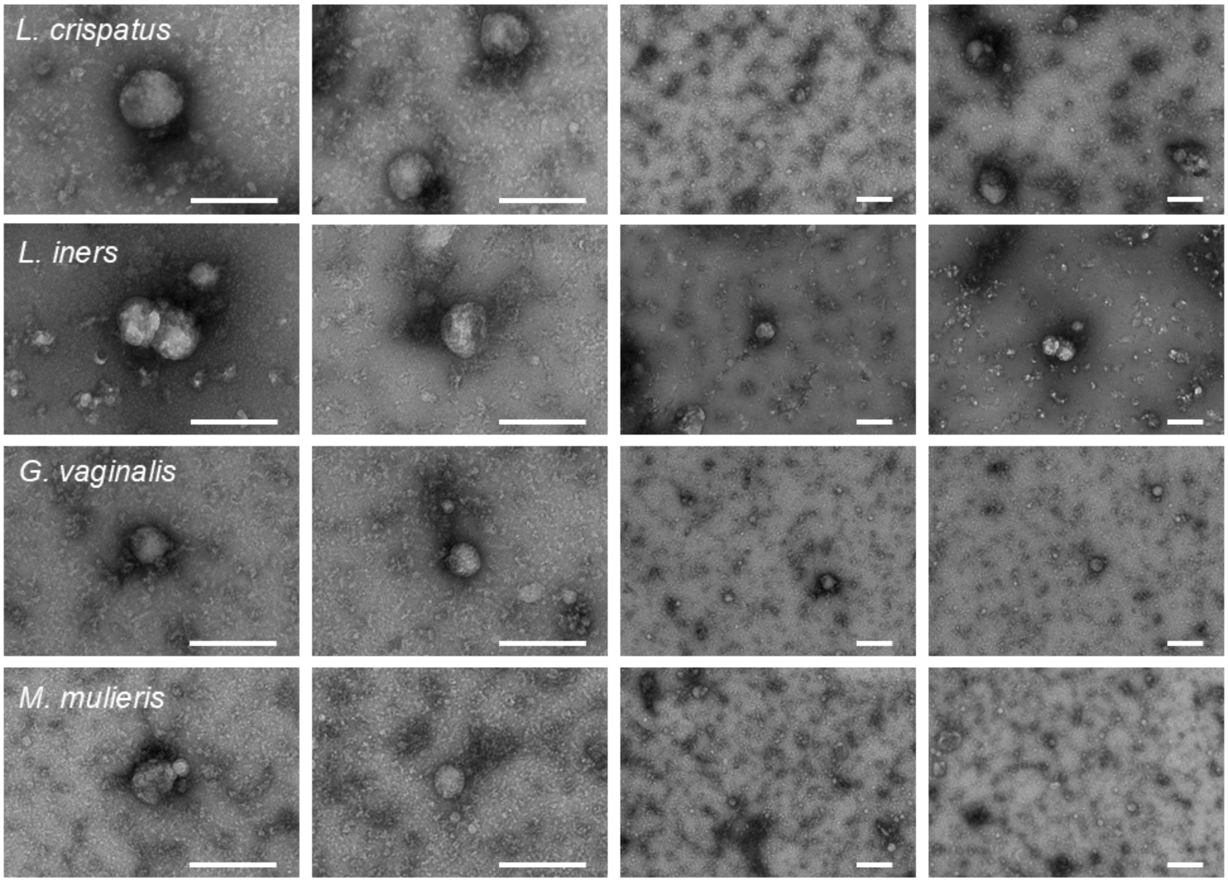


**Supplementary Figure 1: Morphology of isolated bEVs.** Isolated bEVs were confirmed to be membrane-bound particles via TEM. Samples were placed on Formvar/Carbon 200 Mesh grids and negative stained using uranyl acetate. Scale bars denote 200 nm.

**Supplementary Table 1: Cervicovaginal mucus sample characteristics.** Nugent scores and pH for cervicovaginal mucus samples. Nugent scores were determined via Gram staining. Samples were used for multiple-particle tracking experiments within 24 h of sampling.

| SAMPLE | NUGENT | PH |
|---|---|---|
| **1** | 0 | 4.13 |
| **2** | 0 | 4.51 |
| **3** | 2 | 3.62 |
| **4** | 2 | 4.12 |
| **5** | 3 | 4.8 |
| **6** | 6 | 4.28 |
| **7** | 7 | 3.98 |
| **8** | 7 | 5.67 |
| **9** | 8 | 4.7 |
| **10** | 9 | 4.02 |

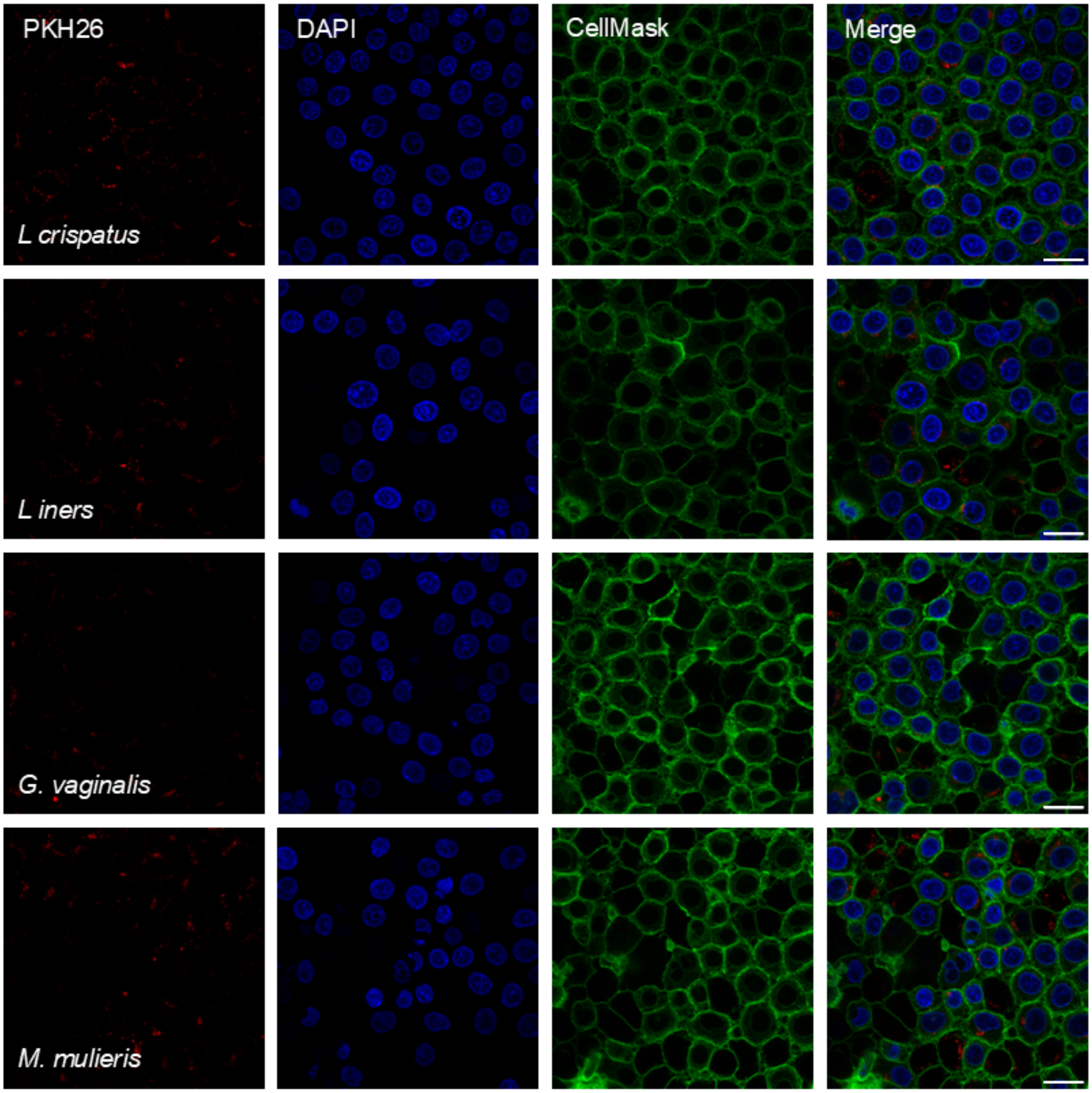


**Supplementary Figure 2: Vaginal epithelial cell uptake of bEVs.** Vaginal epithelial cells demonstrate uptake of bEVs after 24 h. bEVs (PKH26, red) can be seen in the same plane as nuclei (DAPI, blue), within cell membranes (CellMask, green).

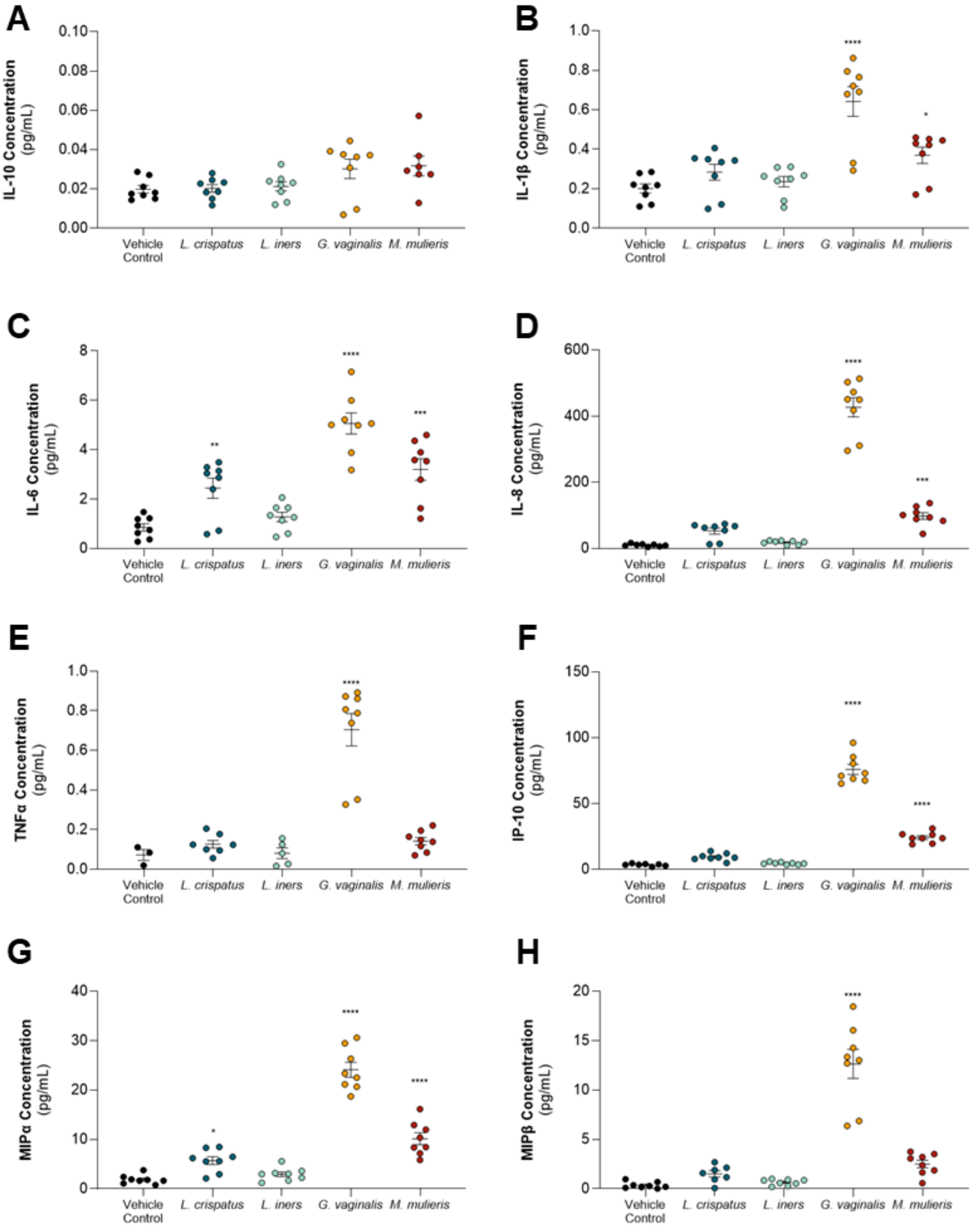


**Supplementary Figure 3: Cytokine concentrations from vaginal epithelial cells after bEV treatment.** *G. vaginalis*- and *M. mulieris*- derived bEVs increase proinflammatory cytokine production in vaginal epithelial cells. (**A**) Treatment with bEVs does not affect vaginal epithelial IL-10 production. (**B**) *G. vaginalis*-derived bEVs (0.64 ± 0.08 pg/ml, $p < 0.0001$), and *M. mulieris*-derived bEVs (0.37 ± 0.04 pg/ml, $p = 0.0425$) increase IL-1β production compared to PBS vehicle control (0.20 ± 0.02 pg/ml) (**C**) *L. crispatus*-derived bEVs (2.44 ± 0.41 pg/ml, $p = 0.0091$), *G. vaginalis*-derived bEVs (5.06 ± 0.43 pg/ml, $p < 0.0001$) and *M. mulieris*-derived bEVs (3.20

± 0.44 pg/ml, $p = 0.0001$) increase IL-6 production compared to PBS vehicle control (0.86 ± 0.15 pg/ml). (**D**) *G. vaginalis*-derived bEVs (426.70 ± 29.04 pg/ml, $p < 0.0001$) and *M. mulieris*-derived bEVs (97.77 ± 10.07 pg/ml, $p = 0.00004$) increase IL-8 production compared to PBS vehicle control (9.53 ± 1.37 pg/ml) (**E**) *G. vaginalis*-derived bEVs (0.70 ± 0.08 pg/ml) increases TNFα production compared to PBS vehicle control (0.07 ± 0.03 pg/ml, $p < 0.0001$). (**F**) *G. vaginalis*-derived bEVs (75.99 ± 3.75 pg/ml, $p < 0.0001$), and *M. mulieris*-derived bEVs (24.22 ± 1.38 pg/ml, $p < 0.0001$) increase IP-10 production compared to PBS vehicle control (3.47 ± 0.34 pg/ml). (**G**) *L. crispatus*-derived bEVs (5.72 ± 0.80 pg/ml, $p = 0.0294$), *G. vaginalis*-derived bEVs (24.22 ± 1.52 pg/ml, $p < 0.0001$) and *M. mulieris*-derived bEVs (10.16 ± 1.20 pg/ml, $p < 0.0001$) increase MIPα production compared to PBS vehicle control (1.89 ± 0.32 pg/ml). (**H**) *G. vaginalis*-derived bEVs (12.6 ± 1.47 pg/ml) increases MIPβ production compared to PBS vehicle control (0.36 ± 0.11 pg/ml, $p < 0.0001$).

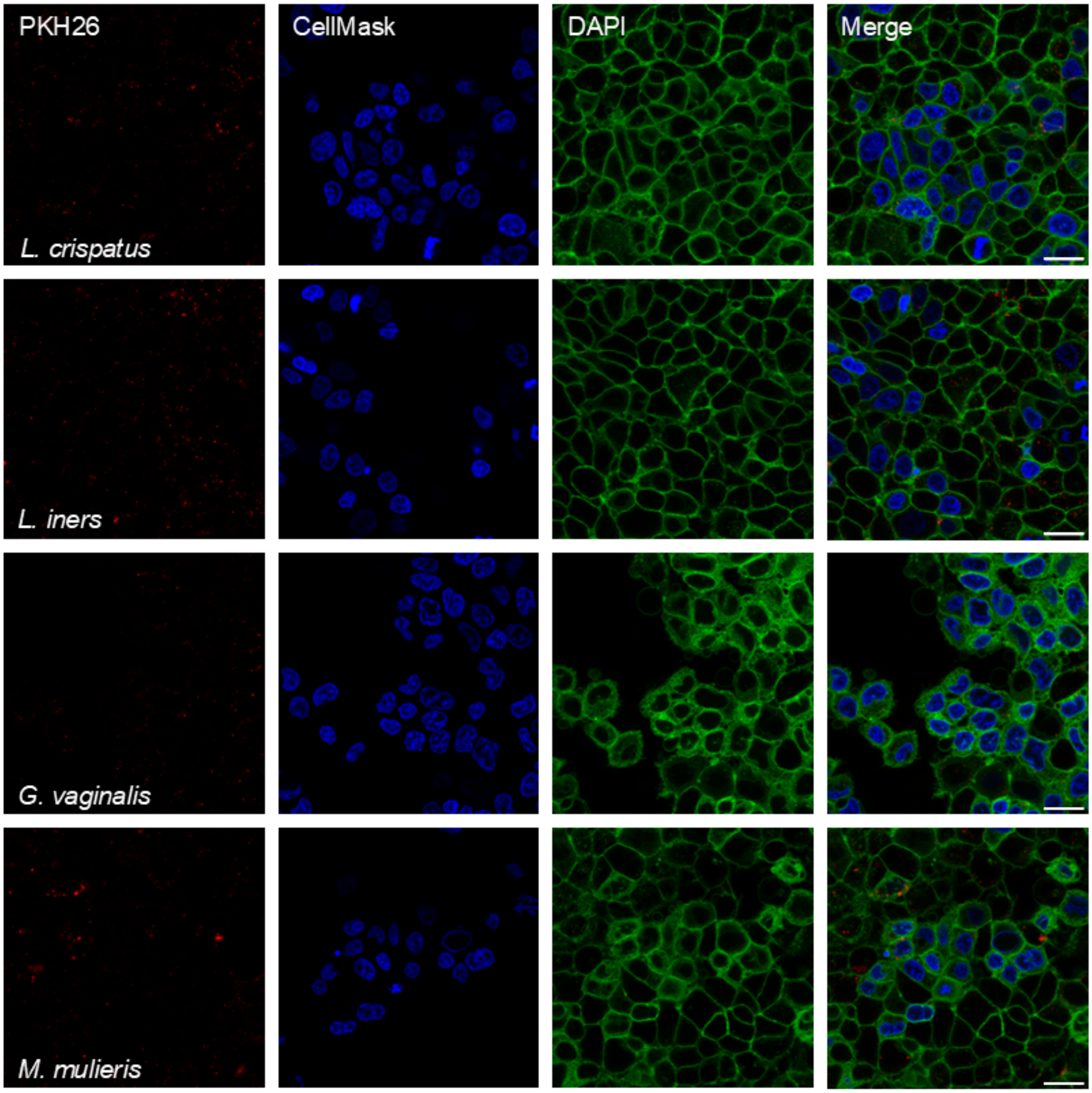


**Supplementary Figure 4: Endometrial cell uptake of bEVs.** Endometrial cells show uptake of bEVs after 24 h. bEVs (PKH26, red) can be seen in the same plane as nuclei (DAPI, blue), within cell membranes (CellMask, green).

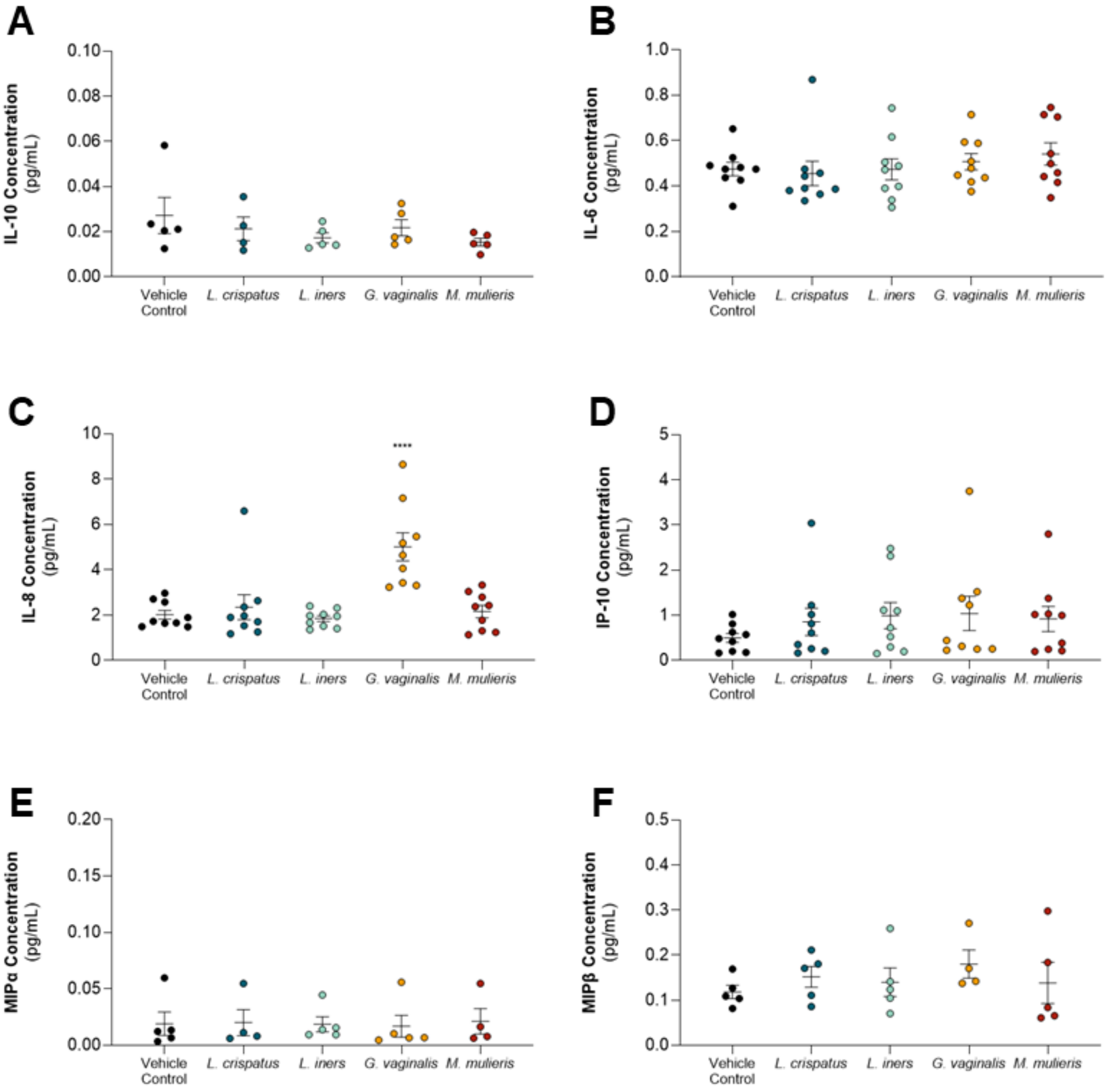


**Supplementary Figure 5: Cytokine concentrations from endometrial cells after bEV treatment** *G. vaginalis*- and *M. mulieris*-derived bEVs increase proinflammatory cytokine production in endometrial cells. (**A**) Treatment with bEVs does not affect endometrial IL-10 production. (**B**) Treatment with bEVs does not affect endometrial IL-6 production. (**C**) *G. vaginalis*-derived bEVs (5.02 ± 0.62 pg/ml) increases IL-8 production compared to PBS vehicle control (2.02 ± 0.19 pg/ml, $p < 0.0001$). (**D**) Treatment with bEVs does not affect endometrial IP-10 production. (**E**) Treatment with bEVs does not affect endometrial MIPα production. (**F**) Treatment with bEVs does not affect endometrial MIPβ production.

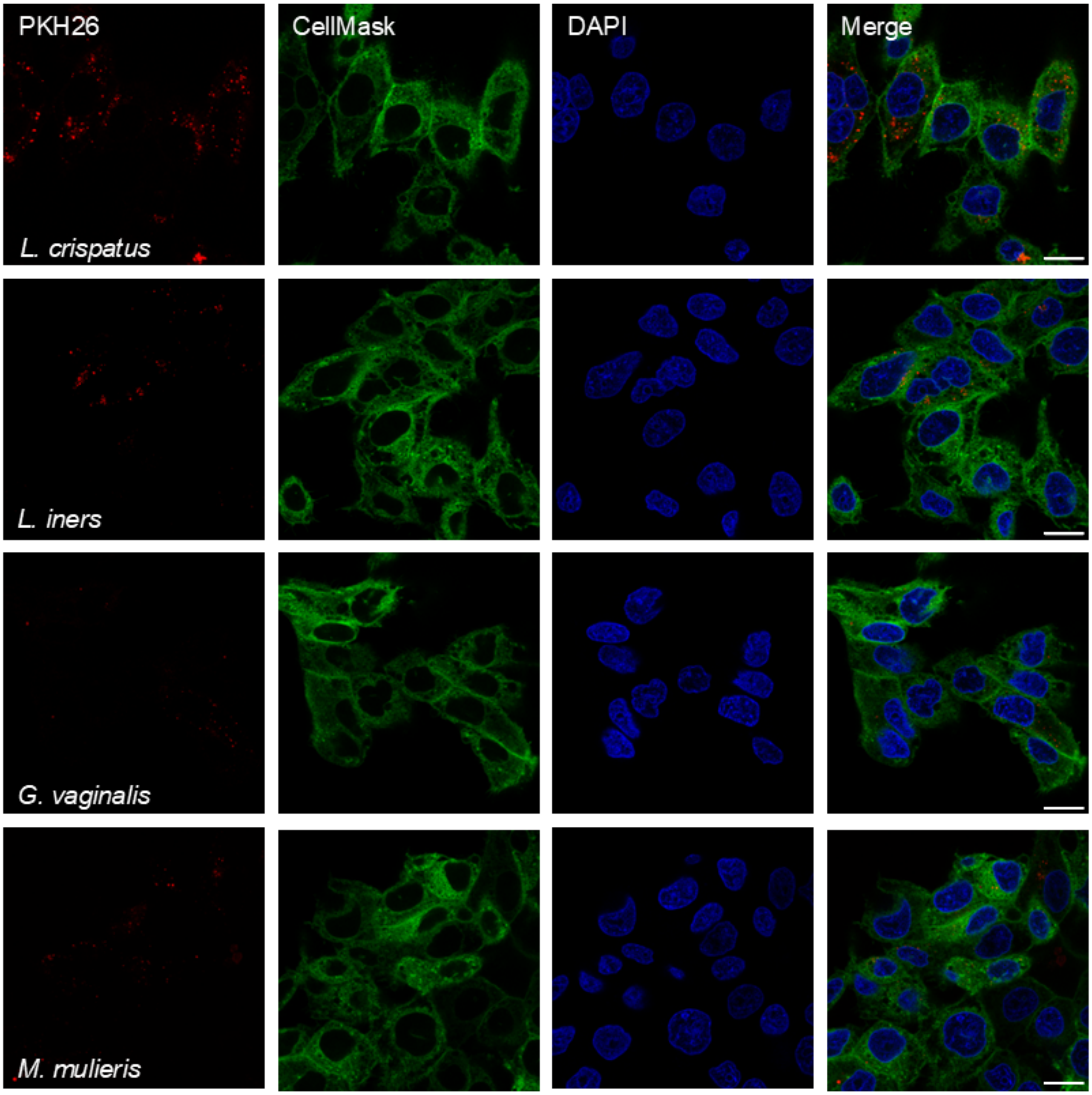


**Supplementary Figure 6**: **Placental cell uptake of bEVs.** Placental cells show uptake of bEVs after 24 h. bEVs (PKH26, red) can be seen in the same plane as nuclei (DAPI, blue), within cell membranes (CellMask, green).

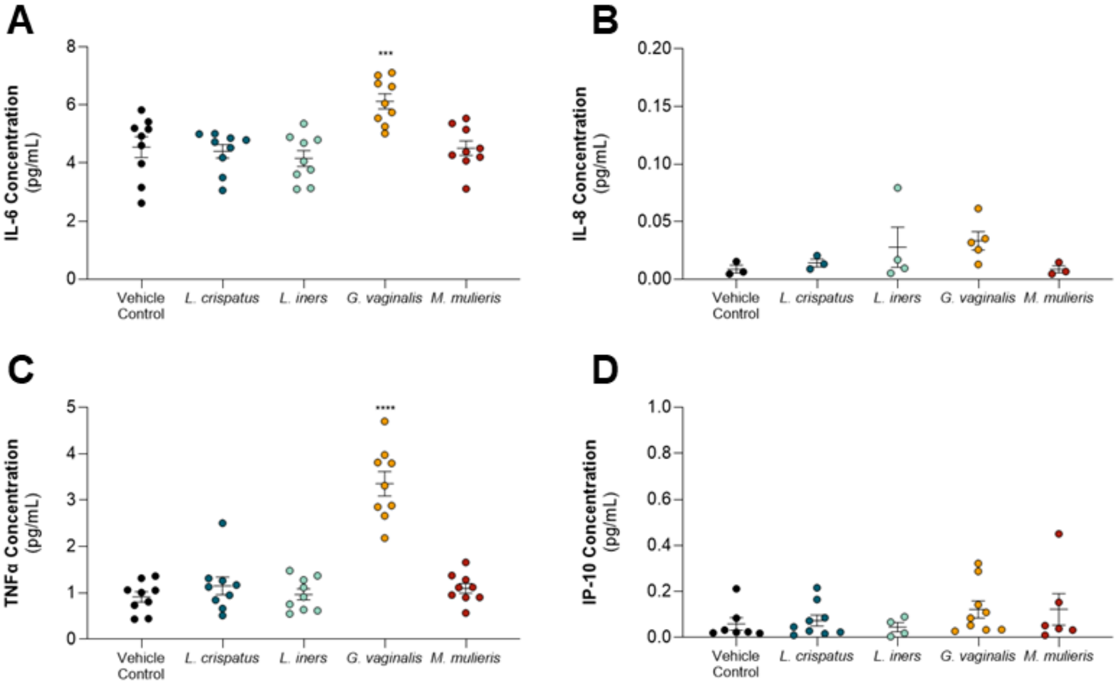


**Supplementary Figure 7: Cytokine concentrations from placental cells after bEV treatment** *G. vaginalis*- and *M. mulieris*- derived bEVs increase proinflammatory cytokine production in trophoblast cells. (**A**) *G. vaginalis*-derived bEVs (6.13 ± 0.27 pg/ml) increases IL-6 production compared to PBS vehicle control (4.52 ± 0.25 pg/ml, $p = 0.0010$). (**B**) Treatment with bEVs does not affect trophoblast IL-8 production. (**C**) *G. vaginalis*-derived bEVs (3.36 ± 0.26 pg/ml) increases TNFα production compared to PBS vehicle control (0.92 ± 0.11 pg/ml, $p < 0.0001$). (**D**) Treatment with bEVs does not affect trophoblast IP-10 production.
